# The use of Statin-class compounds to suppress methanogenesis in lake sediment inoculated microbial fuel cells

**DOI:** 10.1101/2022.08.24.505145

**Authors:** Chamindu Jayathilake, Gayani P. Dilangani, Sampath Bandara, Zumaira Nazeer, Nirath Thilini, Wijendra Bandara, Ajith C. Herath, Godfrey Kyazze, Eustace Y. Fernando

**Affiliations:** Department of Biology, Faculty of Applied Sciences, Rajarata University, Mihintale, Sri Lanka. 50300; Faculty of Technology, Rajarata University, Sri Lanka. 50300; Department of Physics, Faculty of Science, University of Peradeniya, Sri Lanka; Department of Chemistry, Faculty of Applied Sciences, Rajarata University, Sri Lanka. 50300; Faculty of Life Sciences, University of Westminster, 115, New Cavendish Street, London, UK, W1W 6UW

**Keywords:** Statins, Simvastatin, Atorvastatin, microbial fuel cell, methanogenic archaea, power density

## Abstract

Methanogenesis conducted by archaea acts as a competing metabolic pathway that diverts available carbon and electrons away from exo-electrogenic metabolism in mixed culture inoculated microbial fuel cells (MFCs). Statin-class compounds are known to selectively inhibit eukaryotic and archaeal versions of HMG Co-A reductase (class-I) enzyme and the bacterial version of the same enzyme (class-II) is known to be unresponsive to statins. The results of this study demonstrated that the two model statin compounds Simvastatin and Atorvastatin were effective in suppressing methanogenesis in MFCs when applied in moderate concentrations (5 mg/L and 40 mg/L respectively) in MFC anodes. Power densities increased 2 fold compared to control (to 63 ± 1.8 mW/m^2^) and 2.5 fold (to 69.5 ± 1.8 mW/m^2^) with Simvastatin and Atorvastatin addition respectively. There was an almost complete suppression of CH_4_ production with the addition of both statins into MFC anodes as shown by gas composition analysis. Quantitative FISH (qFISH) analysis showed that methanogens *Methanosarcina, Metanobacteria* and *Methanomicrobiales* together with all archaea were almost completely suppressed when statins were supplemented into MFC anodes. This study demonstrated that the statins addition can be used to boost power densities in MFCs.

## 1.0. Introduction

Microbial fuel cells (MFCs) are bioelectrochemical devices that directly convert the chemical energy of organic substrates into electrical energy. Conventional two-chamber type MFCs consist of an anode and a cathode compartment, an external circuit and a cation/proton permeable membrane separating the anode and the cathode. The anode compartment houses the microbial monoculture or the mixed microbial community that catalyzes the key electron transfer reactions that are vital for the function of the MFC system. The cathode compartment is usually abiotic utilizing a noble metal catalyst to drive the oxygen reduction reaction (ORR). However, there are instances where recent research utilized biological cathodes to drive the ORR. Microbes in MFCs transfer electrons on to the anode electrode via exo-electrogenesis, and these electrons travel to the cathode via an external circuit. In addition to electrons, protons are also generated during substrate oxidation in the anode by the exoelectrogens in the anode compartment. These travel through a proton exchange membrane in to the cathode where both the electrons and protons are used up in the reduction of oxygen. The electrochemically active microbial community residing in the anode compartment can rely on many different types of substrates ranging from simple organic compounds such as acetate and glucose or complex organic substrate types such as cellulose, starch, organic waste material or molasses (Chae et al., 2009 and Ullah et al., 2020).

Exo-electrogenic bacteria residing within the anodes of MFCs are vital in driving the electrochemical reactions for biogenic electricity production. Exo-electrogens are characterized by their unique ability to transfer their metabolic electrons into insoluble terminal acceptors that outside their cellular membranes. In their natural habitats, these insoluble terminal electron acceptors are most often iron and manganese oxides. In MFC environments, the anode electrode acts as the terminal electron acceptor for extracellular electron transfer (EET) microbial metabolism. In MFCs, this is done using various methods utilizing different proteins in the process. Two main types have been observed, direct contact EET and Indirect contact EET. In direct EET, there’re two subtypes, 1) electron transfer using direct contact between outer membrane cytochromes and external electron acceptor and 2) electron transfer using bacterial Nano wires known as conductive pili. In indirect EET, microbes utilize mediators, which shuttles electrons from the exoelectrogens on to the external electron acceptor. There are both natural and artificial mediators. Natural mediators (e.g. flavins and pyocyanin) are either produced by the exoelectrogens or an external source (eg. Humic acids) whereas, artificial mediators (e.g. neutral red) need to be added exogenously to MFCs (Logan., 2010).

Inoculum sources for MFCs could vary from monocultures such as *Geobacter* spp., *Shewanella oneidensis, Pseudomonas aeruginosa* and many other bacterial isolates to mixed microbial consortia such as wastewater, activated sludge, anaerobic digester sludge, marine sediments and freshwater sediments (eg. lake and river sediment bacterial communities). The inoculation of mixed microbial consortia in MFCs is generally considered beneficial over monoculture inoculated MFCs due to factors such as the ease of operation, simplicity, lower cost, versatility in substrate requirements and the superior electrochemical performance in most instances (Fernando et al., 2013, Prathiba et al., 2022 and Yaqoob et al., 2021). Although the use of mixed cultures is beneficial in MFC performance and operation, it also brings about several challenges that ultimately results in some level of degradation in

MFC electrochemical performance due to competing metabolic pathways found in mixed bacterial communities other than EET metabolism. Two such competing metabolic fates of available substrates for exo-electrogenic metabolism are fermentative pathways, infiltration of molecular oxygen and methanogenesis. Out of these, fermentative pathways can be alleviated to a large extent by making most of the alternative electron acceptors unavailable in the anode compartments of MFCs (Fernando et al., 2014). Longer-term continuous operation of MFCs ultimately results in exhaustion of many of the alternative electron acceptors such as oxygen, sulfate, nitrate and long-chain fatty acids, leading to a marked reduction of fermentative metabolism in MFC anodes. When MFCs are operated using complex carbon sources however, fermentative metabolism is vital in breaking down complex carbon molecules to simpler ones such as acetate that can be directly used in biogenic electricity generation (Koók et al., 2021). The formation of acetate and acetyl Co-A (acetogenesis) from the breakdown of complex organic substrates in the MFC anodes then promotes exo-electrogenic metabolism where, metabolic electrons yielding from acetate are taken-up by the MFC anode electrodes.

The other major competing metabolic pathway to microbial exo-electrogenesis is methanogenesis pathways conducted by archaea (Chae et al., 2010). Especially, in a mixed microbial consortium where methanogens occupy a prominent place, acetoclastic methanogenesis can be expected to use-up a significant proportion of the acetate content available for exo-electrogenesis in MFC anodes. Therefore, significant levels of methanogenesis taking place within MFC anodes becomes a significant drain on available carbon for exo-electrogenesis and can significantly degrade the electrochemical performance in MFCs. Therefore, the primary goal of this study was to use known Statin-class methanogen inhibitor compounds in order to restrict and suppress the action of methanogens in MFC systems.

Statins are a class of fungal derived compounds which inhibits 3-hydroxy-3-methylglutaryl coenzyme A (HMG Co-A) reductase enzyme, thereby, disrupting the mevalonate pathway. It’s a commonly used drug for lowering low density lipids (LDL) levels plasma (Boucher et al., 2001). It prevents the conversion of HMG Co-A in to mevalonate, which is the precursor for cholesterol synthesis. Mevalonate function as precursor for many other non-steroidal isoprenoidic compounds as well (Ábrego-Gacía et al., 2021). This allows them to be a multifunctional drug with a plethora of physiological effects. Because of this multifunctional nature of statins, they’re able to exert their effect toward both Eukaryotes and Prokaryotes, especially on most archaea including methanogens.

HMG Co-A reductase in archaea, has an essential role in the formation of archaeal cell membrane. It is required in the synthesis of isoprenoid ethers which is the main component of archaeal cell membranes. The lipid composition of the archaea consists of chains of isoprenoids linked to the sn-glycerol-1-phosphate backbone by ether bonds. Synthesis of complex isoprenoid ether lipids is inhibited by the inhibition of the mevalonate synthesis. So the inhibition of mevalonate synthesis by statins through the inhibition of HMG Co A reductase class 1 enzyme leads to the inhibition of archaeal cell membrane synthesis (Ábrego-Gacía et al., 2021). In turn, this suppresses the archaeal community, resulting in suppression of methanogenesis. Since aceticlastic methanogens are prevalent in MFC anodes inoculated with lake sediment, they use up acetate which otherwise would have been utilized by exoelectrogens for electricity generation, thereby reducing the efficiency of the microbial fuel cell. Therefore, it can be expected that selective suppression of such methanogenic archaea communities in mixed-culture inoculated MFCs could yield increased power densities.

In this study Atorvastatin and simvastatin were selected as the candidate compounds to study the potential suppression of methanogens in MFC anodes.

## 2.0. Materials and methods

### 2.1. Construction of two-chamber microbial fuel cells

The two-chamber MFCs and its connected gas collection apparatus was setup as shown in Figure – 1. Three identical MFC units with gas collection units were used and one identical additional unit was used as the control. A 5×5 cm woven carbon microfiber material electrode was used as the Anode. The carbon microfiber material was obtained from Carbon Energy WT (Taiwan) as custom fabricated woven carbon material for electrode construction. The carbon microfiber material had an average fiber diameter of 8 ± 0.5 µm as per the manufacturer data. Each woven carbon microfiber electrode was approximately 4 mm thick. A 5×5 cm carbon micro fiber material electrode pre-treated with KOH according to Wang et al., 2013 in order to functionalize it and was used as the cathode. No other noble metal catalyst such as platinum was used to catalyse the oxygen reduction reaction (ORR) in the cathode. Nafion 212 membrane (Membranes International, USA) was used as the proton exchange membrane. Nafion 212 membrane was pre-treated as per the manufacturer instructions prior to installation in MFCs. Anode and cathode compartments had a volume of 250ml. A 50 mM, pH 7.1 phosphate buffered minimal media as described earlier (Nazeer and Fernando., 2022) was used as the anolyte. A 50mM, pH 7.1 phosphate buffer solution was used as the catholyte.

**Figure-1:**
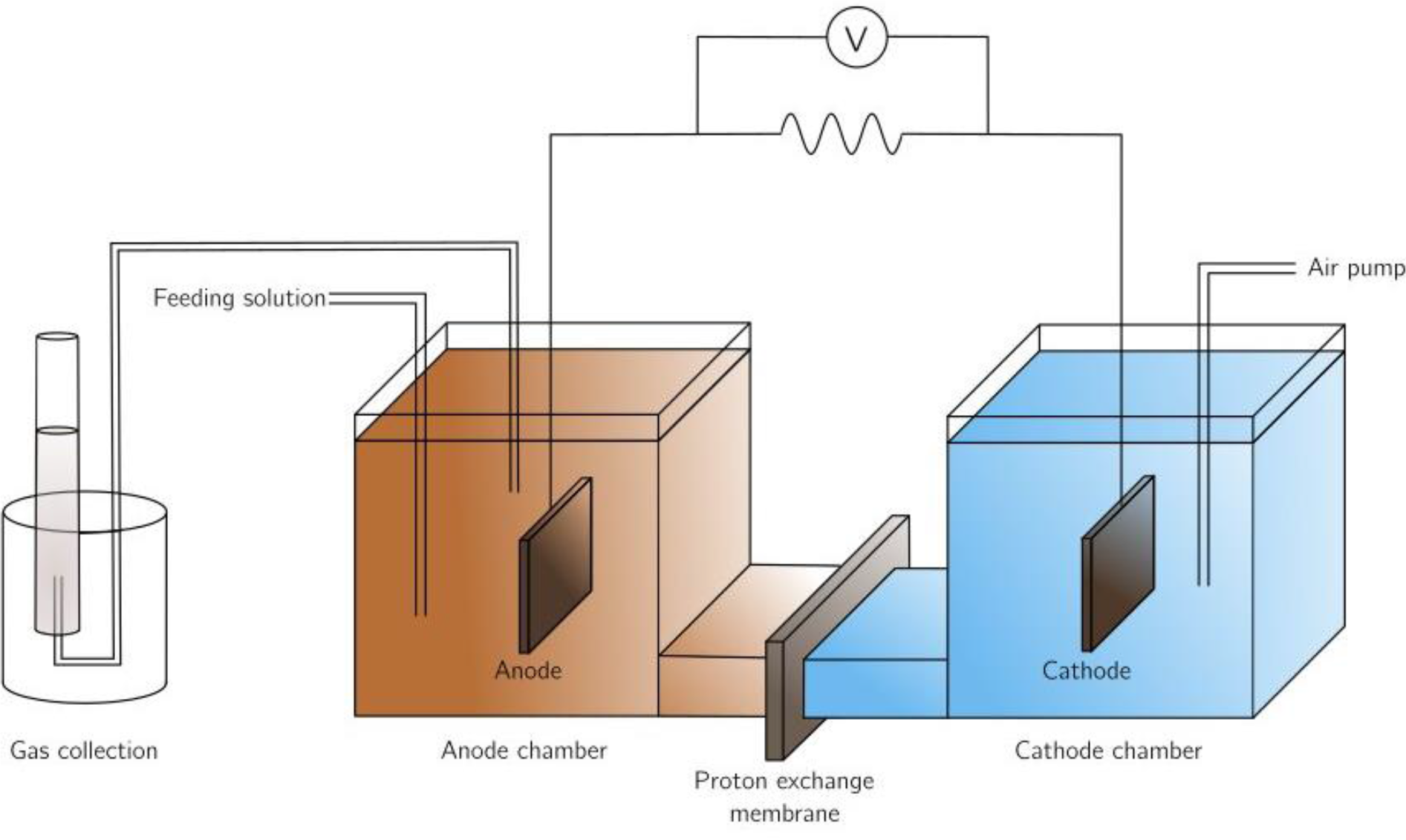
Schematic diagram of the components, the external gas collection apparatus and the external circuit of the customized two-chambered MFC system used in this study

The catholyte of the MFC was continually aerated using an air pump and an air-stone sparger at an aeration rate of 200 mL/min.

### 2.2. Lake sediment sample collection and MFC inoculation

All MFCs were inoculated using anaerobic lake sediment from Mihintale Lake (8°21’47.6” N 80°30’32.8” E). The sediment from the depth of about 10 cm from the sediment surface was collected in to 50 mL tubes under minimal exposure to air. Lake sediment sludge was inoculated into the anolyte minimal medium described earlier in section 2.1. After inoculation of the anode chamber the anode chamber was sealed using silicone sealant to make it airtight. Then an air pump was connected to the cathode to aerate the cathode. The inoculated sediment indicated pyrite mineral deposition as well as rich methanogenic activity. MFCs were inoculated with approximately 10 g of lake sediment. Once inoculated, the anode chambers were fully sealed-off using a silicone sealant except for the feeding port and gas collection port. Glucose was added to the anolye at a concentration of 10 mM was supplemented and replenished as necessary as the substrate for MFCs.

### 2.3. Gas collection during MFC experiments

Gases produced and collected within the headspace of MFCs was collected in a separate vessel by upward displacement of air as shown in Fig-1. Gas volumes produced per MFC anode were recorded as a daily (24 hours sampling interval) recording. All of the control experiments throughout the duration of the study exhibited a baseline daily average gas production rate of 106 ± 10.4 ml/day.

### 2.4. Statin compound supplementation experiments for suppression of methanogenesis in MFCs

Two statin class compounds Simvastatin and Atorvastatin were tested for methanogen suppression activity in MFCs. After MFCs have stabilized to produce a stable voltage profile, statin class compounds were supplemented in an incremental concentration series together with glucose substrate.

Atorvastatin was added in a concentration series of 1 mg/L, 5mg/L, 10mg/L, 20mg/L and 40mg/L. Simvastatin was added in the concentration series 0.5mg/L, 1.0mg/L, 2.0mg/L, 4.0mg/L, 5.0 mg/L and 6.0mg/L. Each statin addition experiment in MFCs was initiated from fresh MFCs acclimated with only lake sediment inoculum that were not previously used in statin addition experiments. All experiments were conducted in duplicates and each experiment included a control experiment containing lake sediment as the inoculum without the addition of any statin class compounds.

### 2.5. Electrochemical assessments of MFCs

An autonomous data logging system (PicoLog™ 1012, Pico Technology, UK) was connected parallel to the 600 Ω external resistance and the voltage across the external resistance (R_ext_) was continually measured and recorded at an interval of 30 seconds. The external circuit connecting the anode and cathode electrodes was connected through a 600Ω resistor. This was used for the continued logging of data at a data recording interval of 30 seconds for the duration of experiments. The circuit was opened when necessary to connect a R_ext_ range from 22 Ω to 1 MΩ to conduct measurements needed to construct polarization curves and power-current plots. Performance characterization of MFCs using polarization curves and power-current plots was conducted as described earlier (Nazeer and Fernando., 2022). The Coulombic efficiency estimations of the MFCs during Statin compound addition experiments were conducted as described earlier in Liu et al., 2005.

### 2.5. Determination of Coulombic efficiency of MFCs during statin additions

Coulombic efficiency (CE%) during the operation of MFC systems during statin addition experiments was determined as described earlier in previous studies (Fernando et al., 2012 and Logan et al., 2006).

Percent Coulombic efficiency was calculated as per the following expression;

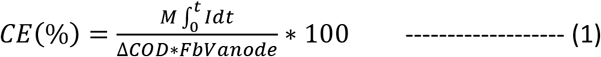

Where, M is the molecular weight of Oxygen (M_r_ = 32), I is the current (A), F is the Faraday’s constant (96,485 C mol^-1^), b is the number of electrons exchanged per mol of oxygen (4), V_anode_ is the working volume of the anode (L) and ΔCOD is the change of COD over time (t) (g COD L^-1^). The chemical oxygen demand (COD) determinations for CE% determinations were conducted as earlier described using the closed-reflux titrimetric method (Fernando et al., 2012 and Fernando., 2014)

### 2.6. Quantitative fluorescent in-situ hybridization (qFISH) for quantification of methanogenic populations and activity

Quantitative fluorescent in-situ hybridization (qFISH) was used to quantitatively study the population abundances of archaea in general and selected methanogenic communities in particular during statin compound additions into lake sediment inoculated MFCs. Samples from MFC anode control experiments and statins added experiments were drawn for qFISH analysis and they were immediately fixed in paraformaldehyde (PFA) as described earlier (Fernando et al., 2019). Archaea populations in general and selected methanogenic archaea populations were quantified using FISH probes as described in earlier work (Dawson et al., 2012). FISH probes ARCH915 (5’ -GTG CTC CCC CGC CAA TTC CT 3’) targeting most archaea, MS821 (5’-CGC CAT GCC TGA CAC CTA GCG AGC -3’) targeting *Methanosarcina*, MB311 (5’ - ACC TTG TCT CAG GTT CCA TCT CC -3’) targeting *Methanobacteria*, MG1200b (5’ -ACC TTG TCT CAG GTT CCA TCT CC -3’) targeting *Methanomicrobiales* were used. The general EUBmix probe set (EUB 338, EUB 338 II, and EUB 338 III) (Daims et al., 1999) targeting all bacteria were used obtain relative quantitative FISH signal intensities for selected methanogenic archaeal members in the community from the total population of bacteria and archaea. Fluorescence microscopy was conducted using an Olympus BX61 fluorescence microscopy system (Olympus, Japan). FISH probes were labelled with 5(6)-carboxyfluorescein-N-hydroxysuccinimide ester (FLUOS) or with the sulfoindocyanine dyes (Cy3) at the 5’ end of the probes (Sigma Aldrich company, Germany). FISH sample preparation was conducted as described earlier (Fernando et al., 2019) and qFISH analyses were conducted as described in a previous study (Daims et al., 2006). Quantitative FISH analyses were based on 30 random fields of view taken at ×630 magnification on each sample. Relative signal intensities for each FISH probe used was calculated using ImageJ and Daime (Daimes et al., 2006b) image analysis software.

### 2.7. Qualitative and quantitative headspace gas analysis during MFC experiments

Head space gas (approximately 10 mL) of microbial fuel cells was collected after 12 hours from each feeding in to vacuum tubes. The composition and relative quantity (as a percentage of the total) of the collected gas was analyzed. Composition and relative quantity of the headspace gases (CH_4_, CO_2_, H_2_S and O_2_) during MFC experiments were measured using a GasBoard 3200 Plus handheld biogas analyzer (Cubic-Ruiyi Instrument Co., Ltd, China).

### 2.8. Scanning electron microscopy (SEM) and Fourier transform infrared spectroscopic (FTIR) analysis of functionalized microfiber electrode surfaces

Woven electrode Microfiber samples were imaged for SEM on a Carl Zeiss EVO 18 (Germany) scanning electron microscope with optical magnification range of 20–135 ×, electron magnification range of 5x - 1,000,000 x, maximal digital zoom of 12 ×, acceleration voltage of 10 kV, equipped with secondary electron (SE) and energy dispersive X-ray spectrometer (EDS) detectors, with nominal resolution of 10 nm or less. The microscope had a temperature controlled sample holder (temperature range −25°C to 50°C).

Chemical functionalization occurring on the base pretreated cathodes was analyzed using FTIR as described earlier in Dissanayake et al., 2021. Briefly, dried cathode microfiber material pieces were cut out from base pretreated electrode MFC experiments and were analyzed using FTIR spectrometer (Shamadzu, UK, National Institute for Fundamental Studies, Sri Lanka). Fully dried samples were placed on the instrument analysis pedestal and spectra were obtained in the mid-IR region (400 cm^-1^ – 4000 cm^-1^) in the transmittance mode.

### 2.9. Statistical analysis

All experimental data plotted in graphs are the means of duplicate experiments and the error bars represent the standard error of the mean (SEM). Statistical analysis of data was conducted by one way analysis of variance (ANOVA) using Prism GraphPad 5.0

## 3.0. Results and discussion

### 3.1. MFC electrochemical performance during statin compound addition experiments

Supplementation of Simvastatin into MFC anodes exhibited a clear concentration – dependent increase of maximum obtainable power densities from the MFC systems (Figure 2A). Approximately, a two-fold increase in maximum power densities over the control experiments (no Simvastatin – 31.5 ± 1.2 mW/m^2^) could be achieved when MFC anodes were supplemented with 5 mg/L of Simvastatin (63.1 ± 1.8 mW/m^2^). This represents a significant improvement of power density over the control experiment (P<0.05) and a power density fold increase of more than 2 when simvastatin is introduced at a concentration of 5 mg/L into the MFC anode. Similarly, additions of Atorvastatin into the MFC anode produced a concentration-dependent increase in power densities up-to an Atorvastatin concentration of 40 mg/L. The maximum power density recorded in Atorvastatin experiments was 69.5 ± 1.9 mW/m^2^ (Figures 2C and 2D). This increase in power density after the addition of Atorvastatin was a significant increase in power density over the control experiment (P<0.05) and the fold increase of power density over the control experiments was in excess of a factor of 2.5. The increase in power densities in MFC anodes was incremental and dependent upon the addition of Simvastatin up-to a concentration of 5 mg/L. Further additions of Simvastatin at 6 mg/L and beyond indicated a reduction in maximum achievable power densities from the MFC system (reducing to 57.9 ± 5.7 mW/m^2^ at Simvastatin concentration of 6 mg/L) (Figure 2B). This could probably be due to any cytotoxic effects of Simvastatin to the exo-electrogenic bacterial community when introduced at a higher concentration. This effect was also observed in experiments of Atorvastatin additions where, supplementation of Atorvastatin at concentrations exceeding 40 mg/L produced a reduction in power densities (Figure 3B). The highest power density obtainable by addition of Atorvastatin was 69.5 ± 1.9 mW/^2^, at the Atorvastatin concentration 40 mg/L. Further additions reduced the P_max_ values (Figures 2C and 2D). It was also evident that further additions of statins (> 5 mg/L for simvastatin and > 40 mg/L for Atorvastatin) did not produce any significant increases in Coulombic efficiencies as well (Figure 3). These observation are suggestive of the two statin compounds tested becoming cytotoxic to the anode exoelectrogenic bacterial community after a threshold concentration (5 mg/L for Simvastatin and 40 mg/L for Atorvastatin additions).

**Figure – 2:**
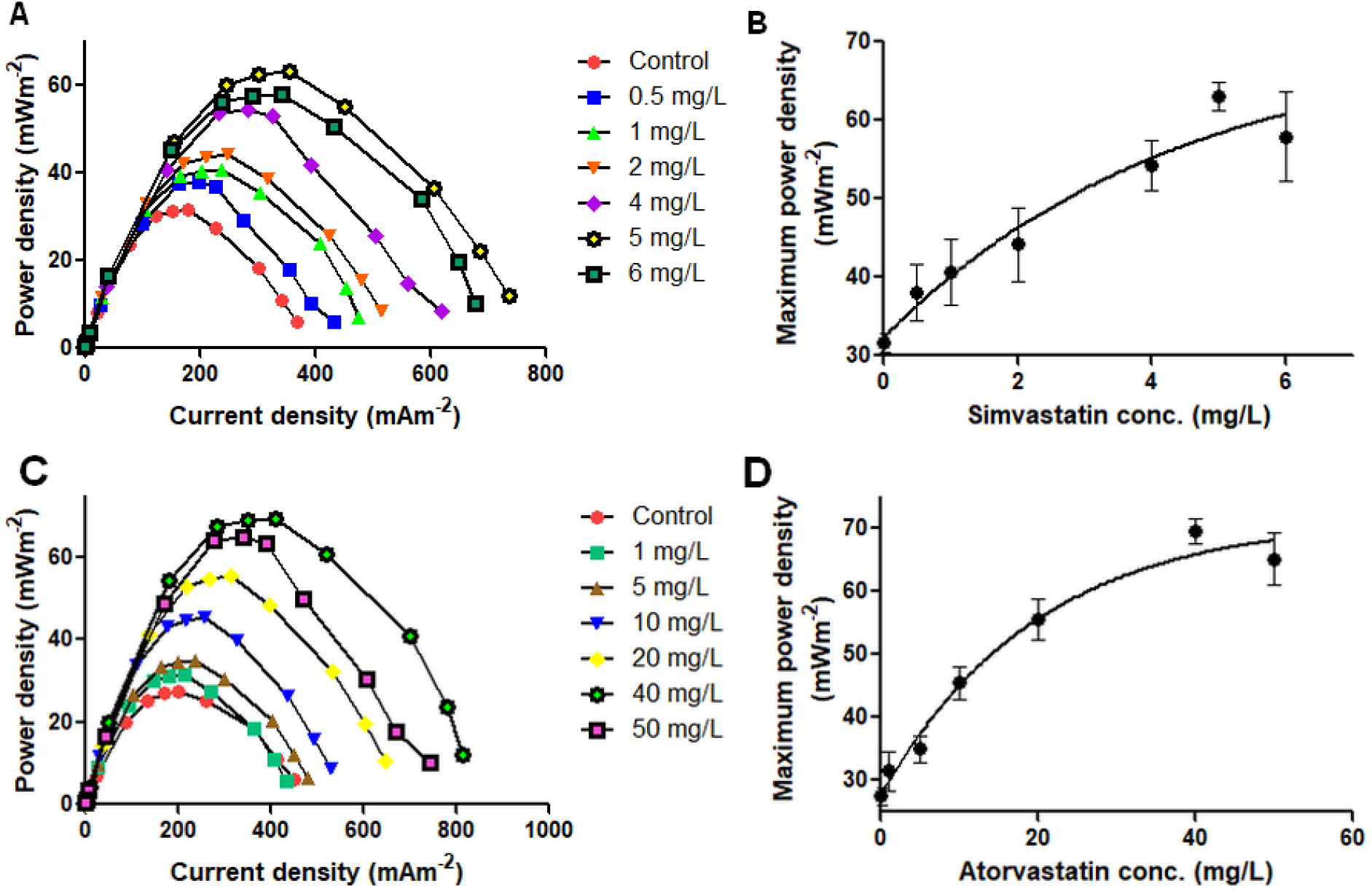
(A) Current-power plots of MFC experiments utilizing incremental concentrations of Simvastatin (B) the effect of incremental supplementation of Simvastatin on peak power densities (P_max_) produced in MFCs (C) Current-power plots of MFC experiments utilizing incremental concentrations of Atorvastatin (D) the effect of incremental supplementation of Atorvastatin on peak power densities (P_max_) produced in MFCs.

**Figure-3:**
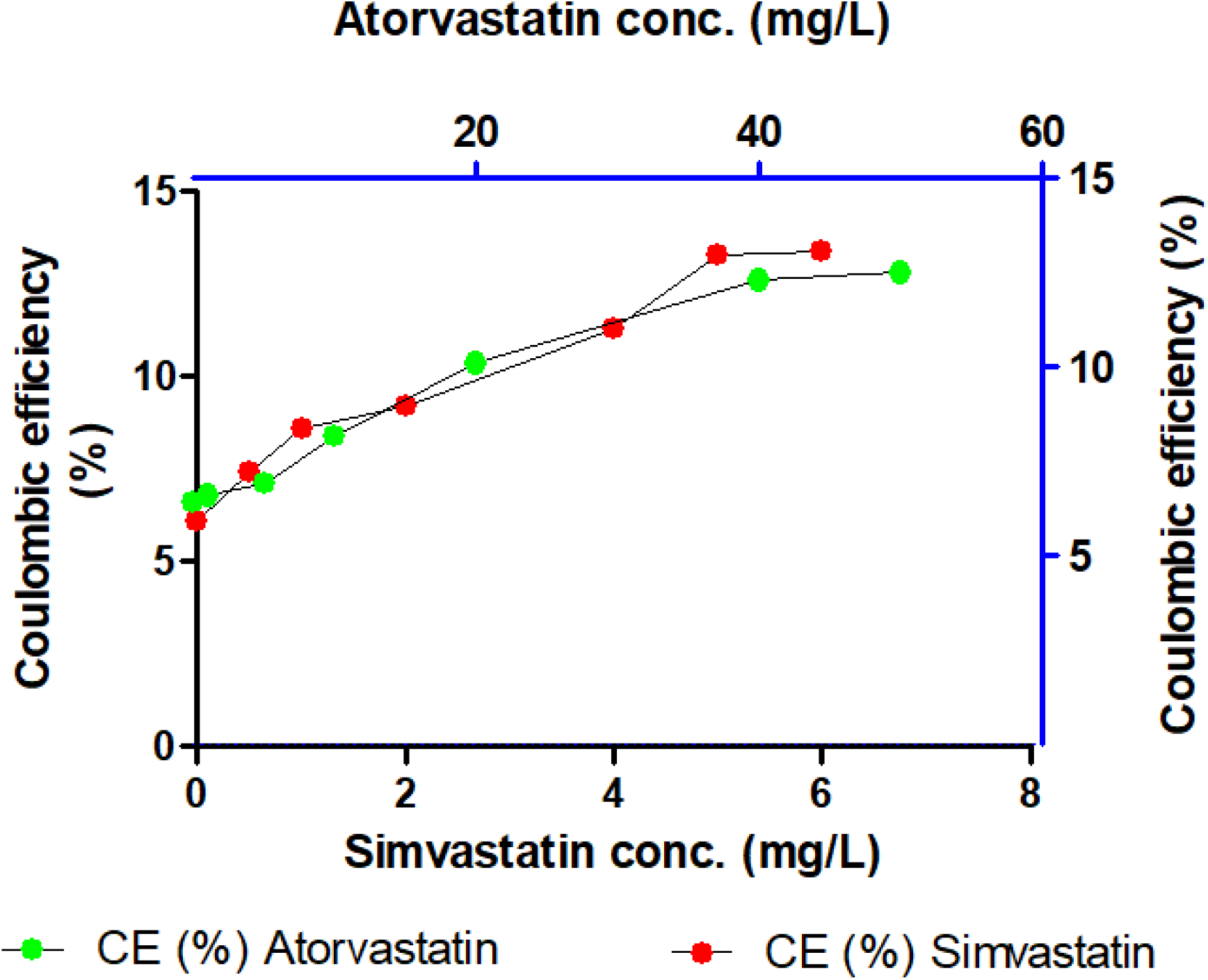
The effect of statins supplementation (Simvastatin and Atorvastatin) in incremental concentrations on Coulombic efficiency (CE %) of MFCs

Earlier studies conducted by Graziano et al., 2015 on medically-important bacterial monocultures demonstrated that Simvastatin has a minimum inhibitory concentration (MIC) of 15.65 mg/L for *Staphylococcus aureus* 29213. Also another study has shown that Atorvastatin, Simvastatin and Rosuvastatin exert antibacterial effects on Methicillin-sensitive *Staphylococcus aureus* (MSSA), methicillin-resistant *Staphylococcus aureus* (MRSA), vancomycin-susceptible *Enterococci* (VSE), vancomycin-resistant *Enterococcus* (VRE), *Acinetobacter baumannii, Staphylococcus epidermidis*, and *Enterobacter aerogenes* (Masadeh et al.,2012). The inhibitory effects of the two statin compounds on exoelectrogenic bacteria in MFC anodes in this study could be caused by the previously reported antibacterial effects of these statin compounds at higher concentrations.

### 3.2. Improvement of Coulombic efficiencies in MFCs by statin additions

Coulombic efficiencies (CE) of MFCs increased more than two fold compared to the control experiments during the additions of both statin class compounds into MFC anodes. The increasing trend of CE during additions of Simvastatin and Atorvastatin was concentration-dependent. CE improved from 6.4% to 12.4% when Atorvastatin concentration was raised from 0 mg/L (control experiment) to 50 mg/L. Similarly, the CE increased from 6.1% (control experiment) to 13.4% when Simvastatin concentration was raised to 5 mg/L in the MFC anodes (Figure 3). The increase in CE is further evidence for increased availability of carbon and as a result, increased availability of electrons from carbon substrates to be channeled into exo-electrogenic bacterial metabolism. Suppression of other competing pathways such as methanogenesis will free-up carbon sources and the molar equivalents of electrons that can be released from the extra carbon. Similar enhancements of CE by blocking fermentative metabolic pathways in MFC anodes have been demonstrated in several previous studies (Kim et al., 2011 and Ebadinezhad et al., 2019).

The gradual and concentration-dependent increase in CE with the supplementation of statins is strong evidence to suggest that supplementation of statins into MFC anodes in moderate concentrations is an effective strategy to circumvent the channeling of available carbon and their metabolic electrons into competing metabolic pathways such as methanogenesis. Furthermore, statins supplementation into MFC anodes can channel metabolic electrons predominantly into exo-electrogenic metabolism and therefore increase CE and maximum power densities (P_max_) of MFC systems. Coupling the findings of the current study with blocking of other competing pathways such as fermentative metabolism of microbial communities may allow further enhancements to CE and P_max_ outputs of various MFC systems.

### 3.3. Reduction of methanogenesis by the addition of statins into MFC anodes

The supplementation of both statin class compounds into MFC anodes made significant reductions of total gas volumes produced and their methane contents compared to the control experiments where no statin additions were made. Total gas volumes produced per day were nearly halved when both Atorvastatin and Simvastatin were supplemented in moderate concentrations (6 mg/L for Simvastatin and 50 mg/L for Atorvastatin respectively) (Figures 4A and 4C). The reduction in total gas volume was dependent upon the concentration of the two supplemented statin compounds.

**Figure-4:**
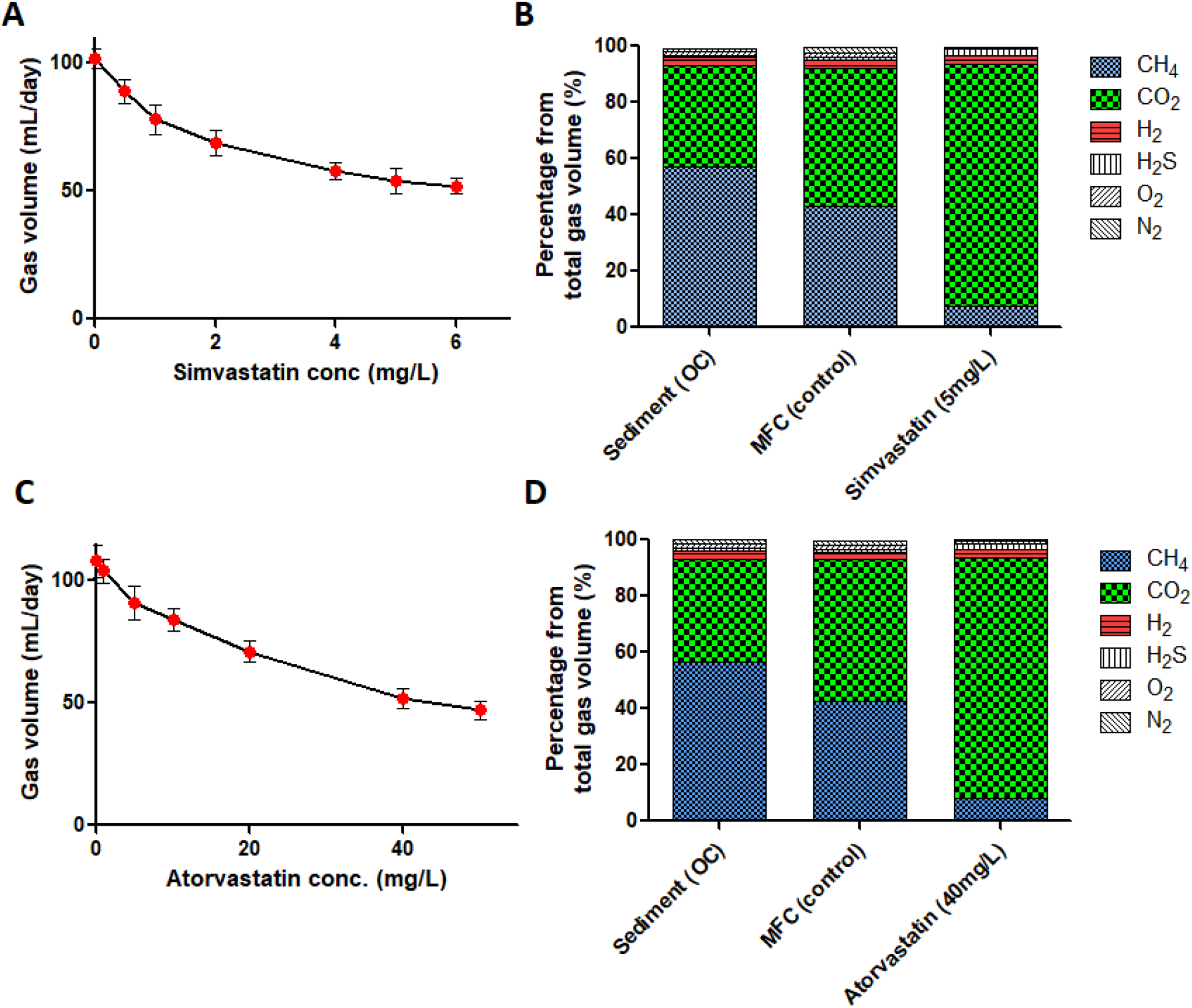
(A) The effect of incremental addition of Simvastatin on gas production in MFC anodes (B) the changes to collected gas composition in presence of Simvastatin (C) The effect of incremental addition of Atorvastatin on gas production in MFC anodes (D) the changes to collected gas composition in presence of Atorvastatin * OC – MFC operated at open-circuit configuration (infinite external resistance) to mimic lake sediment environment.

The gas composition analysis data further suggested that the methane content of the collected gas was significantly lower in statins supplemented MFC anodes compared to the control experiments.

Additions of Atorvastatin into MFC anodes at increasing concentrations demonstrated a concentration-dependent suppression of total gas production and methanogenesis (Figures 4C & 4D) (Supplementary table S2). Total gas production nearly halved (52 ± 3.9 mL/day) and the volume fraction of methane was about four times less at 7.9% CH_4_ at Atorvastatin concentrations of 40 mg/L compared to the control experiments (42.7 % CH_4_) where no Atorvastatin was added.

Similarly, supplementation of Simvastatin into MFC anodes indicated a clear suppression of total gas volume produced and noticeable drop in methanogenesis. A concentration – dependent decrease of total gas volume produced per day in MFC anodes when the Simvastatin concentration was raised (Figure 4B). Methanogenesis was suppressed to as little as 7.6% of the total gas composition when the Simvastatin concentration was raised to 5 mg/L over the control experiment (no Simvastatin) where, the volume fraction of methane in the collected gas was 43.2% (Figure 4B and Supplementary table S1).

These observations clearly indicate that methanogenesis during statin addition experiments (both Simvastatin and Atorvastatin) was severely suppressed. The dominant gas by volume fraction at the highest statin additions was CO_2_ (Figures 4B & 4D). This clearly suggests that the carbon provided by the substrate additions are channeled towards exo-electrogenic metabolism rather than entering methanogenic pathways under the addition of statin class compounds. Methanogenesis is a metabolic capability held exclusively by the prokaryotic organism group archaea in the biosphere. Therefore, the reduction of biogenic CH_4_ gas content and CH_4_ percentage in the produced headspace gas in this study is direct evidence of suppression of methanogenesis. It is also direct evidence to suggest that the methanogen populations in MFC anodes supplemented with statin compounds were selectively suppressed and other prokaryotic organisms such as bacteria in general and exo-electrogenic bacteria in particular, were unaffected by addition of statins. Several other earlier studies investigated the possibility of suppression of methanogens in MFCs by a range of methods including introducing stress conditions for the microbial community (Chen et al., 2010) and making changes to the external resistances of the MFC external circuits (Jung and Regan., 2011). However, none of the previously reported studies successfully attempted the usage of exogenous additions of statin class compounds as methanogen inhibitors of MFC anodes.

### 3.4. Quantitative dynamics of methanogen populations during statins addition experiments in MFCs

During quantitative FISH analysis, the FISH signal intensity signals for the probe representing the total archaea population in the sample (ARCH 915) indicated a significant reduction when statins were introduced into the MFC anodes. This reduction of signal intensity for total archaea population appeared to be concentration-dependent on the exogenous additions of statin compounds. However most of the signal loss for the archaea-specific probe was produced during the initial low concentration additions of statins (Figure 5B). This is clear quantitative evidence to suggest that statins addition into the MFC anodes produces a suppressive effect on the total archaea population compared to the control experiments where no statins were present.

**Figure – 5:**
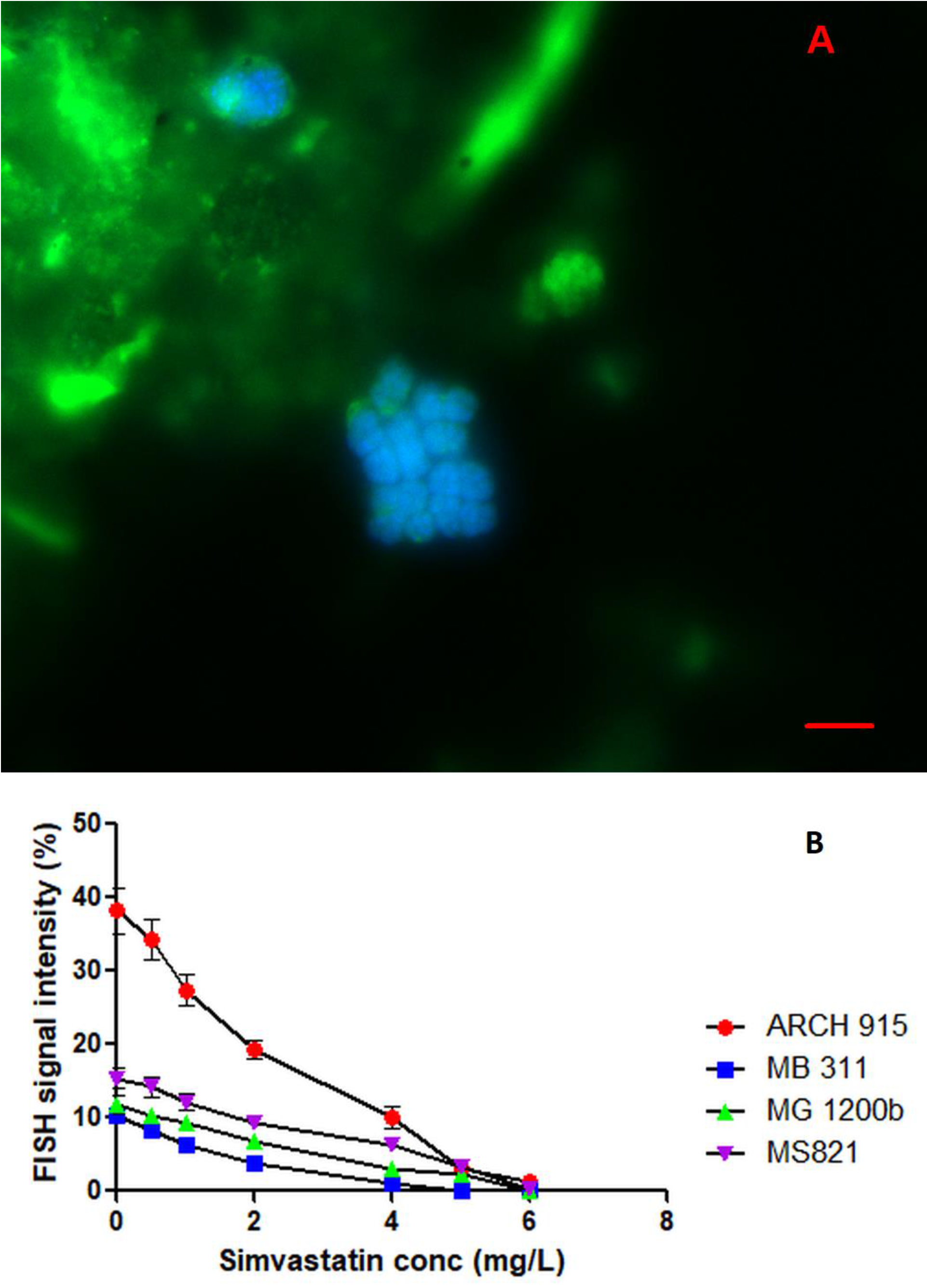
A) Representative qFISH micrograph showing hybridization signals overlap from ARCH915 representing all archaea (Green) and MS821 representing the methanogenic archeaon *Methanosarcina* (Cyan signal produced by the overlap of green and blue FISH signals) (Scale bar = 10 µm). B) Changes to qFISH signal intensities (ARCH 915, MB311, MG1200b and MS821) in the presence of increasing Simvastatin concentration in MFC anodes.

Similarly specific known groups of methanogenic archaea were clearly suppressed during statins addition experiments. Methanogenic archaeal group *Methnosarcina, Methanobacteria* and *Methanomicrobiales* represented by the FISH probes MS821, MG1200b and MB311 (Figure 5A) respectively were all clearly suppressed by the additions of statins into the MFC anode compartments. Methanogenic archaeal groups *Methnosarcina, Methanobacteria* and *Methanomicrobiales* represented by the FISH probes MS821, MG1200b and MB311 were present between 11% - 15.4% in the control MFC anodes where no statins were supplemented (Figure 5B). This suggests significant methanogenic activity by these methanogenic organism groups and it is further corroborated by the gas composition analysis results where, in-excess of 50% of the collected headspace gas was methane. However, with the addition of statins, it was clearly evident that all of the methanogenic archaeal groups represented by the FISH probes MS821, MG1200b and MB311 produced no detectable qFISH signal at the highest concentrations of Simvastatin and Atorvastatin when they were supplemented into the MFC anodes (Figure 5B). The complete absence of qFISH signal for the three groups of methanogenic archaea *Methnosarcina, Methanobacteria* and *Methanomicrobiales* is a clear indication that methanogenic activity was suppressed to a large extent by the addition of Simvastatin and Atorvastatin at concentrations 5 mg/L and 40 mg/L respectively. At the highest statin concentrations (6mg/L Simvastatin and 50 mg/L Atorvastatin), the qFISH signal was almost completely suppressed for all archaea (ARCH915 FISH probe) at 1.4% of the total qFISH signal. Out of this, qFISH signals for methanogenic archaea was not detectable, suggesting that they were completely eliminated from the MFC anode microbial community by the presence of Statin compounds at high concentations.

Moreover, at this stage, almost the entirety of the qFISH signal (98.6%) was coming from bacteria (represented by the FISH probes EUB 338, EUB 338 II, and EUB 338 III, suggesting that the suppressed archaeal community, including methanogens was replaced by bacteria.

### 3.5. Possible mechanisms of methanogen suppression in MFCs by statins additions

Statins are known to act by inhibiting the activity of the enzyme HMG Co-A reductase. This is done through competing with HMG-Co-A for the binding site. In addition, they alter the conformation of the enzyme after binding. This prevents HMG Co-A reductase from attaining a functional structure (Istvan., 2003). This makes the statin compounds highly effective and specific on HMG Co-A reductase. HMG Co-A reductase catalyzes the conversion of HMG Co-A to mevalonate. This enzyme is found in both eukaryotes and prokaryotes. Through molecular studies and phylogenetic analysis, two classes of HMG Co-A reductases have been identified (Ábrego-Gacía et al., 2021). Class I HMG Co-A reductase enzymes of eukaryotes and some archaea including methanogenic archaea and the Class II HMG Co-A reductase enzymes of eubacteria. The class-I HMG Co-A enzyme (found in eukaryotes and archaea) is found to be highly sensitive to statin class compounds and the Class-II HMG Co-A enzyme (bacterial version) was found to be relatively insensitive to the action of Statins (Ábrego-Gacía et al., 2021 and Boucher et al., 2001). HMG Co-A reductase in Archaea, has an essential role in the formation of archaeal cell membrane. It is required in the synthesis of isoprenoid ethers which is the main component of archaeal cell membranes. The lipid composition of the archaea consists of chains of isoprenoids linked to the sn-glycerol-1-phosphate backbone by ether bonds. Synthesis of complex isoprenoid ether lipids is inhibited by the inhibition of the mevalonate synthesis. So the inhibition of mevalonate synthesis by statins, through the inhibition of HMG Co A reductase class-I enzyme leads to the inhibition of archaeal cell membrane synthesis (Ábrego-Gacía et al., 2021). Which in turn suppresses the archaeal community. Thereby, suppressing methanogenesis.

The selective and concentration-dependent inhibitory effect shown by the application of the two statin compounds Simvastatin and Atorvastatin into MFC anodes in this study is most likely due to the ability of these compounds to selectively inhibit the archeal community, including the methanogenic archaea. Furthermore, the supplementation of statins in moderate concentrations is likely to exert minimal or no adverse effect on the bacterial community including the exo-electrogenic and fermentative bacterial communities of the MFC anodes because their version of HMG Co-A reductase enzyme (Class-II) is largely insensitive to statin class compounds.

### 3.6. The use of chemically functionalized, noble metal-free microfiber electrodes in MFCs

The results of this study further demonstrated that reasonably high power densities can be achieved with the use of chemically functionalized woven carbon microfiber cathodes that are devoid of noble metal catalysts such as platinum powder. Average power densities for all control experiments (with no statins supplementation in the anode) was 28.8 ± 3.3 mW/m^2^ (Figures 2A and 2C). With the supplementation of statins these power densities could be elevated by a factor between 2 – 2.5.

Scanning electron micrographs of the carbon microfiber material indicated that the highly fibrous nature of the woven material is capable of producing a very high surface area to volume ratio (S/V ratio) (Figure 6A). This will likely create a large surface area for ORR to occur in the woven carbon cathode material. Similar effects were reported by earlier studies where, power densities were improved by the use of material having a large S/V ratio in the cathode (Delord et al., 2017 and Feng et al., 2010). Furthermore, it was confirmed by FTIR analysis that the woven carbon microfibers were functionalized with hydroxyl groups following treatment with KOH (Figure 6B). The base treated carbon microfiber material indicate the emergence of many additional major peaks compared to the untreated micro fiber material at 1355cm^-1^, 2848 cm^-1^ and 2915 cm^-1^ wavelengths. The strong peak at 1355cm^-1^ corresponds to –OH bending vibrations of phenol groups. The other two additional peaks at 2848 cm^-1^ and 2915 cm^-1^ corresponds to –OH stretching vibrations. All three of these additional peaks confirms the hydroxyl group surface functionalization of the base treated micro fiber material. Hydroxyl groups functionalization on carbon cathodes was found to be a good replacement ORR catalyst for noble metal ORR catalysts in an earlier study conducted by Wang et al., 2013. These findings imply that cheap MFC electrodes devoid of noble metal ORR catalysts could still be used effectively to obtain higher power densities, if their anode compartments were supplemented with moderate concentrations of statins to suppress competing methanogenic metabolic pathways.

**Figure – 6:**
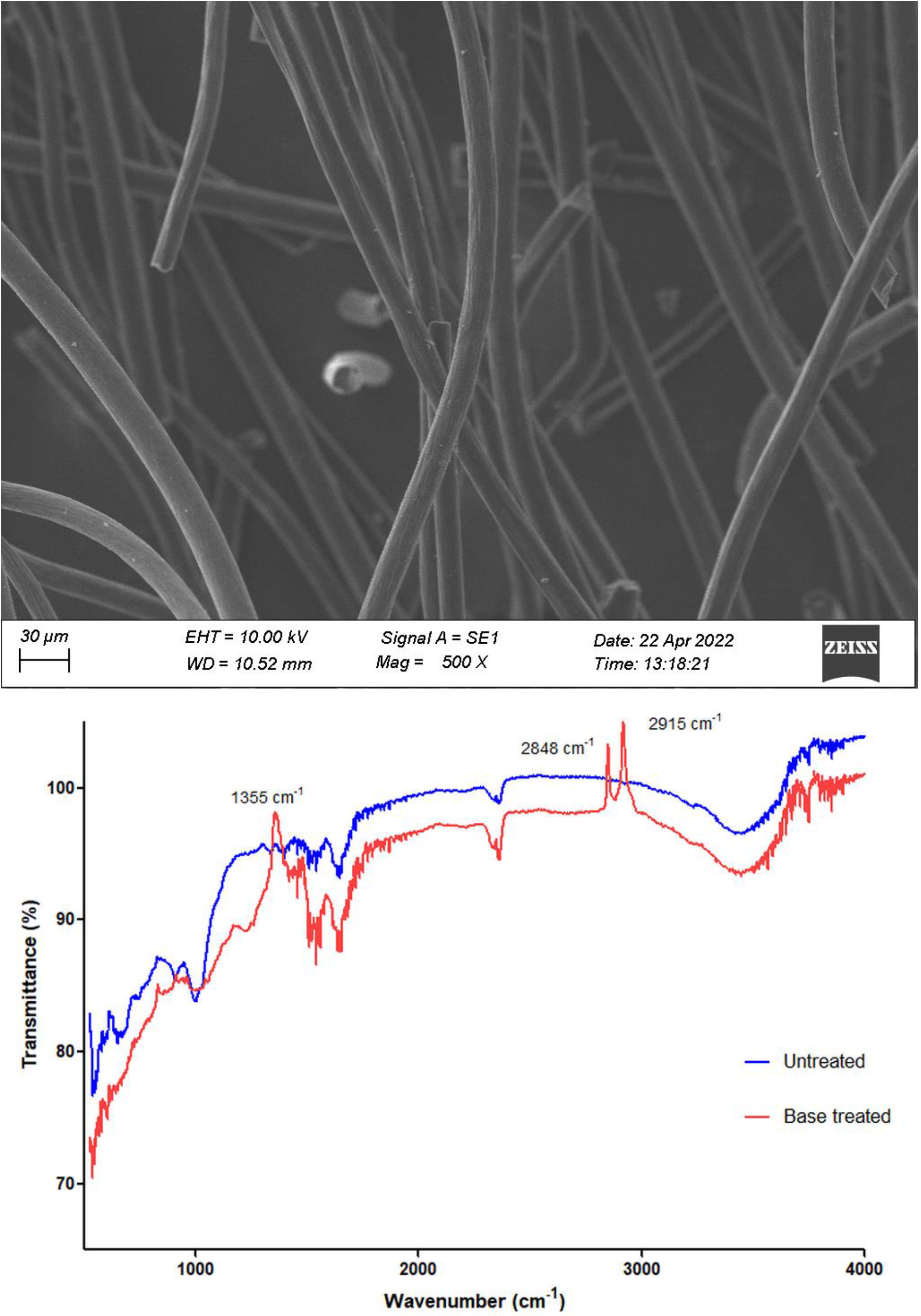
(A) SEM image of KOH treated microfiber cathode material and (B) Overlay of FTIR spectra from untreated carbon microfiber cathode material and functionalized carbon microfiber material using KOH pretreatment

## 4.0. Conclusions

The outcomes of this study has demonstrated for the first time that commonly used statin class compounds Simvastatin and Atorvastatin can be used for selective suppression of methanogenic archaea in lake sediment inoculated MFC anodes. When used in moderate concentrations, (5 mg/L Simvastatin and 40 mg/L Atorvastatin) they could significantly boost the obtainable power densities compared to control experiments (more than 2 fold and 2.5 fold for Simvastatin and Atorvastatin respectively). A noticeable reduction of total gas production and a significant drop in CH_4_ content in the collected gas was also indicative of a large suppression of methanogenic archaea after supplementation of both statins. Direct quantification of hybridization signals for FISH probes designed specifically for archaea further indicated an almost complete suppression of key methanogenic populations in the MFC anodes supplemented with statins. The implications of this work are that statins could be used to effectively be used to boost power densities of mixed-culture MFCs by suppression of methanogenesis.

## 5.0. Acknowledgements

Technical staff who aided the work in various ways at all collaborating institutions and laboratories are thankfully acknowledged.

