## Supplementary data tables for "The use of Statin-class compounds to suppress methanogenesis in lake sediment inoculated microbial fuel cells"

| | $CH_4$ (mL) | $CO_2$ (mL) | $H_2$ (mL) | $H_2S$ (mL) | $O_2$ (mL) | $N_2$ (mL) |
|---|---|---|---|---|---|---|
| **Sediment (OC)** | 57.1 | 36.1 | 2.7 | 0.7 | 1.3 | 1.2 |
| **MFC (control)** | 43.2 | 48.9 | 2.8 | 1.2 | 1.6 | 1.8 |
| **Simvastatin (5mg/L)** | 7.6 | 86.2 | 2.9 | 2.2 | 0.4 | 0.4 |

**Supplementary table S1**: Composition of collected headspace gas during Simvastatin supplementation experiments (Simvastatin concentration at 5 mg/L)

| | $CH_4$ (mL) | $CO_2$ (mL) | $H_2$ (mL) | $H_2S$ (mL) | $O_2$ (mL) | $N_2$ (mL) |
|---|---|---|---|---|---|---|
| **Sediment (OC)** | 56.4 | 36.9 | 3.0 | 0.8 | 1.4 | 1.4 |
| **MFC (control)** | 42.7 | 50.2 | 2.6 | 1.2 | 1.3 | 1.4 |
| **Atorvastatin (40mg/L)** | 7.9 | 85.9 | 2.9 | 2.1 | 0.6 | 0.5 |

**Supplementary table S2**: Composition of collected headspace gas during Atorvastatin supplementation experiments (Atorvastatin concentration at 40 mg/L).
